# Bridging the divide: bacteria synthesizing archaeal membrane lipids

**DOI:** 10.1101/448035

**Authors:** Laura Villanueva, F.A. Bastiaan von Meijenfeldt, Alexander B. Westbye, Ellen C. Hopmans, Bas E. Dutilh, Jaap S. Sinninghe Damsté

**Affiliations:** NIOZ Royal Netherlands Institute for Sea Research, Department of Marine Microbiology and Biogeochemistry, and Utrecht University, P.O. Box 59, 1797AB Den Burg, Texel, The Netherlands.; Theoretical Biology and Bioinformatics, Science for Life, Utrecht University, The Netherlands.; Centre for Molecular and Biomolecular Informatics, Radboud Institute for Molecular Life Sciences, Radboud University Medical Centre, The Netherlands.; Faculty of Geosciences, Department of Earth Sciences Utrecht University., P.O. Box 80.021, 3508 TA Utrecht, The Netherlands.

## Abstract

Archaea synthesize membranes of isoprenoid lipids that are ether-linked to glycerol, while Bacteria/Eukarya produce membranes consisting of ester-bound fatty acids. This dichotomy in membrane lipid composition or ‘lipid divide’ is believed to have arisen after the Last Universal Common Ancestor (LUCA). A leading hypothesis is that LUCA possessed a ‘mixed heterochiral archaeal/bacterial membrane’, however no natural microbial representatives supporting this scenario have been shown to exist today. Here, we demonstrate that bacteria of the Fibrobacteres-Chlorobi-Bacteroidetes (FCB) group superphylum and related candidate phyla encode a complete pathway for archaeal membrane lipid biosynthesis in addition to the bacterial fatty acid membrane pathway. Key genes were expressed in the environment and their recombinant expression in *E. coli* resulted in the formation of a ‘mixed archaeal/bacterial membrane’. Our results support the existence of ‘mixed membranes’ in natural environments and their stability over large evolutionary timescales, thereby bridging a once-thought fundamental divide in biology.

---

Lipid membranes are essential for all cellular life forms to preserve the integrity and individuality of cells, as well as having a direct influence in the maintenance of energy metabolism. Lipid membranes are also key in differentiating the domains of life. Bacteria and eukaryotes have membranes formed by fatty acids linked to glycerol-3-phosphate (G3P) via ester bonds, while archaea have membranes made of isoprenoid alkyl chains linked by ether linkages to glycerol-1-phosphate (G1P), leading to an opposite stereochemistry of the glycerol phosphate backbone^1^. This segregation in lipid membrane composition, or ‘lipid divide’, has been hypothesized to have appeared early in the evolution of microbial life from LUCA, but the nature of the lipid membrane of LUCA and its subsequent differentiation in Bacteria and Archaea remain unknown. Some studies have proposed that a non-cellular LUCA lacked a lipid membrane^2-4^. A recent study suggested that the membrane of LUCA was formed by fatty acids and isoprenoids without the glycerol phosphate backbone as a requirement to have a lower membrane permeability that could sustain a proton gradient^5^. The most parsimonious hypothesis may be that LUCA had a heterochiral lipid membrane composed of both G1P and G3P together with fatty acids and isoprenoids^6, 7^, which later diversified into archaeal and bacterial membranes resulting in the ‘lipid divide’. It was originally proposed that this differentiation may have been driven by heterochiral membrane instability^8, 9^, but heterochiral membranes are in fact stable^10^ and a recent study has proven that they are, in some cases, more robust to environmental stresses^11^.

Another critical issue in the ‘lipid divide’ is the membrane lipid composition of eukaryotes. Multiple lines of evidence indicate that eukaryogenesis encompassed an endosymbiosis event of a bacterial cell into an archaeal host^12-17^. Thus, the bacterial-like composition of contemporary eukaryotic membranes implies that an early eukaryote had its archaeal-like membrane replaced by a bacterial-like one, possibly through a ‘mixed membrane’ intermediate containing both the archaeal membrane lipids with ether-linked isoprenoids to G1P and the bacterial ones with ester-linked fatty acids to G3P. This would imply that bacterial-like membrane molecules arose twice, in bacteria and in eukaryotes. The competing syntrophic hypothesis of eukaryogenesis^18, 19^ proposes that the host of the mitochondrial endosymbiont was a bacterium, avoiding the need for a transitional step from an archaeal to eukaryotic membrane. Some eukaryogenesis models also suggest that the membrane transition was facilitated by intensive bacterial lipid transfer from the endomembrane system^20^. Because most models for the origin of eukaryotes require a membrane transitional step similar to the one expected in the ‘mixed membrane’ scenario for LUCA, it is striking that no remnants or natural microbial representatives with a heterochiral ‘mixed membrane’ have yet been described that would support the viability of such a scenario.

Over the years, the concept of the ‘lipid divide’ has been challenged by the identification of traits thought to be characteristic of archaeal membrane lipids in bacteria and *vice versa*. Firstly, some bacteria (and eukaryotes) produce ether-linked lipids^21-27^. Secondly, the biosynthesis of membrane-spanning lipids, another trait thought to be specific for the archaea, also occurs in bacteria^26-30^. However, no bacteria are known to produce membrane lipids based on isoprenoidal chains or possess the ‘archaeal’ G1P stereochemistry. Thirdly, it has been postulated that archaea produce small amounts of fatty acids^31^ and an almost complete biosynthetic pathway for fatty acid synthesis is encoded in many archaeal genomes^32^. Lastly, two uncultured archaeal groups, the marine Euryarchaeota group II and Lokiarchaeota, contain archaeal lipid biosynthesis genes alongside bacterial-like fatty acid and ester-bond formation genes but lack the capacity to synthesize the G1P backbone via glycerol 1-P-dehydrogenase (G1PDH), while they have the genetic background to produce G3P^33^. However, a recent study has observed that G1P-lipids can be synthesized in the absence of G1PDH^11^, which may suggest that these uncultured archaeal groups could synthesize a membrane with both archaeal and bacterial traits^33^. This observation is exciting as Lokiarchaeota are considered the nearest descendants of the archaeal ancestor leading to eukaryotes. Nevertheless, there is no further evidence that the presence of these genes in those two archaeal groups actually leads to the synthesis of ‘mixed heterochiral membranes’, meaning membranes harboring both G3P-bacterial and G1P-based archaeal membrane lipids. Taken together, these observations expose a ‘lipid divide’ that is not as clear-cut as originally thought. Nonetheless, no natural microbial representatives have ever been found that can synthesize both archaeal and bacterial membrane lipids.

Serendipitously, we discovered a living ‘mixed archaeal/bacterial membrane’ bacterium in the Black Sea, a basin whose euxinic waters may resemble the ancient oceans^34^. Members of the phylum *Candidatus* Cloacimonetes of the FCB group superphylum were highly abundant in the deep waters (>10% of the 16S rRNA gene reads, see Fig. 1a). A genome-centric metagenomics approach of the water column was undertaken to shed light on the physiology of these bacteria and four *Ca.* Cloacimonetes draft genomes with substantial to near completeness and low contamination were assembled (see Supplementary Information, Supplementary Table 1). The high abundance of these metagenome-assembled genomes (MAGs) in the deep waters matched the *Ca.* Cloacimonetes 16S rRNA gene abundance profile (Fig. 1b), and phylogenomic analysis based on 43 concatenated core genes confirmed their taxonomic position (Fig. 1c-d). Analyses of the MAGs revealed the lipid biosynthetic pathways of *Ca.* Cloacimonetes. First, genes of the bacterial fatty acid biosynthetic pathway were detected, including the gene (*gps*) encoding glycerol-3-phosphate dehydrogenase (catalyzing the formation of G3P) (see Supplementary Information, Supplementary Table 2), the acyltransferases responsible for the esterification of fatty acids and G3P, as well as genes coding for enzymes involved in downstream reactions^35^ (see Supplementary Table 2). Hence, *Ca.* Cloacimonetes harbors the genes enabling the formation of a normal bacterial membrane.

**Figure 1.**
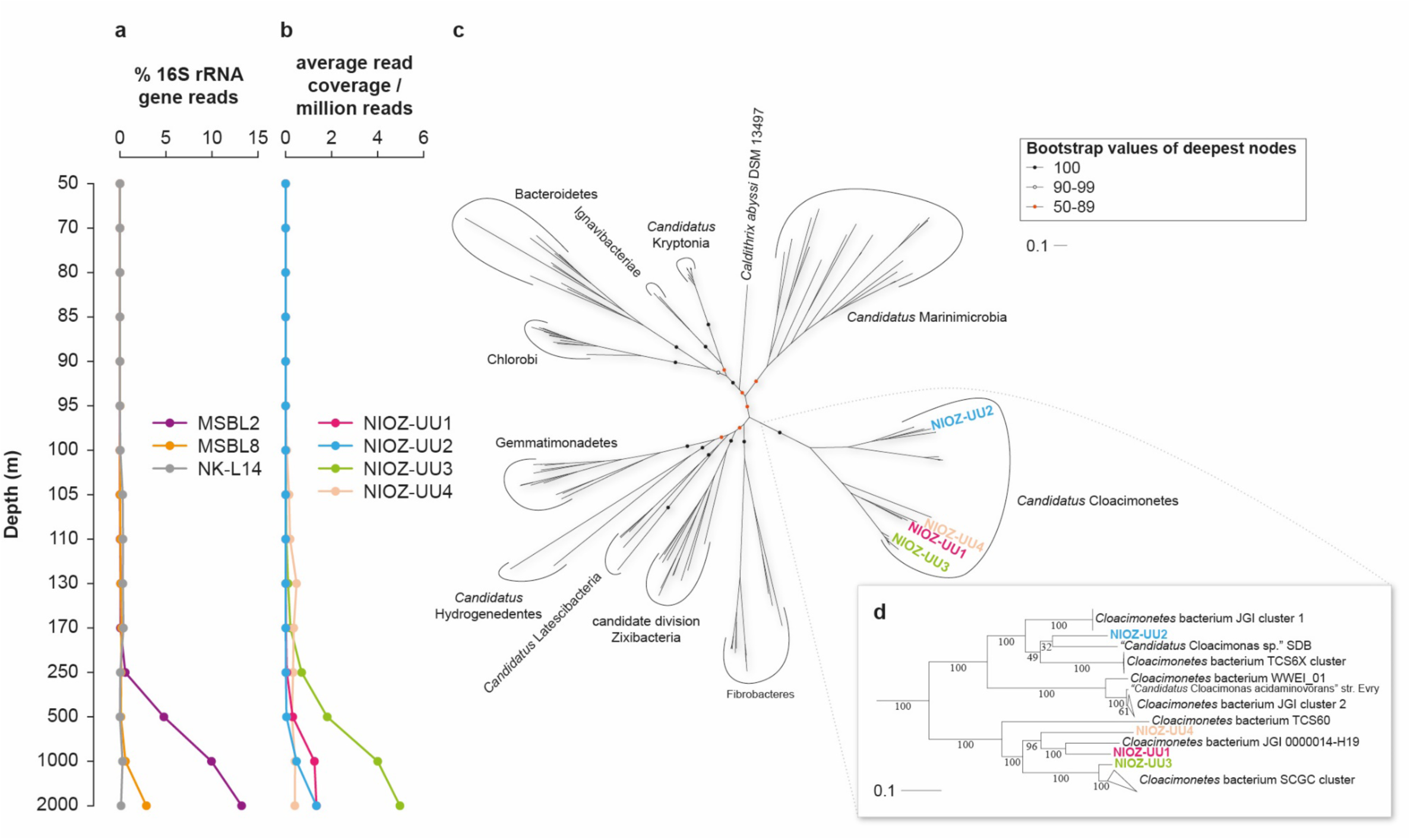
Distribution within the Black Sea water column and phylogeny of *Ca.* Cloacimonetes. **a**, percentage of 16S rRNA gene reads attributed to different *Ca.* Cloacimonetes groups. **b**, Estimated abundance of the *Ca.* Cloacimonetes MAGs. **c**, phylogenetic tree of the FCB group superphylum. **d**, zoom in on the *Ca.* Cloacimonetes phylogeny.

Strikingly, however, putative genes of the archaeal lipid biosynthetic pathway were also detected in all four *Ca.* Cloacimonetes MAGs. A homolog of the archaeal generanylgeranylglyceryl phosphate (GGGP) synthase is co-localized (i.e. encoded in close proximity) in the genomes with an archaeal (*S*)-2,3-di-O-geranylgeranylglyceryl phosphate (DGGGP) synthase (Supplementary Information). These two enzymes together mediate the formation of the two ether bonds between isoprenoid alcohols and G1P resulting in the production of unsaturated archaeol^1^. Distant homologs of both GGGP and DGGGP synthase genes have previously been reported in bacterial genomes, however their presence has never been associated with the production of archaeal lipids. Whereas GGGP synthase activity has been confirmed in only a few bacteria^36^, DGGGP synthase belongs to a large superfamily of UbiA prenyltransferases with several different potential functions^32^, and bacterial homologs were assumed to have divergent activity. However, the close proximity of the two genes in the *Ca.* Cloacimonetes MAGs and their relatively high sequence similarity to archaeal homologs led us to hypothesize that this bacterium encodes GGGP and DGGGP synthase activity and is thus capable of synthesizing archaeal-like membrane lipids.

The possibility of a chimeric composition of the MAGs resulting in a spurious detection of GGGP and DGGGP synthase genes in these organisms was discarded by both *in silico* analyses and by experimentally amplifying and resequencing one of the scaffolds containing these genes from the original Black Sea water samples (Supplementary Information). The presence of *Ca.* Cloacimonetes GGGP and DGGGP synthase gene transcripts in the Black Sea water confirmed that they were also expressed (Supplementary Information). Previous studies have determined that GGGP synthase homologs of the phylum Bacteroidetes have *in vitro* GGGP synthase activity on G1P like the archaeal GGGP synthase^36^, rather than the heptaprenyl synthase activity detected in the bacterial PcrB orthologs detected in *Bacillus subtilis*^37^. To biochemically verify the predicted enzymatic activity in *Ca.* Cloacimonetes, we recombinantly produced its GGGP synthase protein in *E. coli*. The purified protein was found to catalyze formation of GGGP from geranylgeranyl diphosphate (GGPP) in an enzymatic assay (Fig. 2a). Then we tested if the *Ca.* Cloacimonetes GGGP and DGGGP synthases could support the formation of a ‘mixed membrane’ in a bacterial cell, by co-expressing them in an *E. coli* optimized for production of GGPP and G1P, the likely substrates of the two enzymes (see Methods). Cells producing both GGGP and DGGGP synthase contained significant amounts of phosphatidylglycerol archaeol with 8 double bonds (Fig. 2b), the expected intermediate in the biosynthesis of archaeal membrane lipids in the absence of a specific geranylgeranyl reductase in *E. coli*^38, 39^. Hence, our experiments provide strong evidence for potential synthesis of archaeal lipids by this group of bacteria, the final validation depending on the isolation of these elusive bacteria from their environment, such as from the deep waters of the Black Sea.

**Figure 2.**
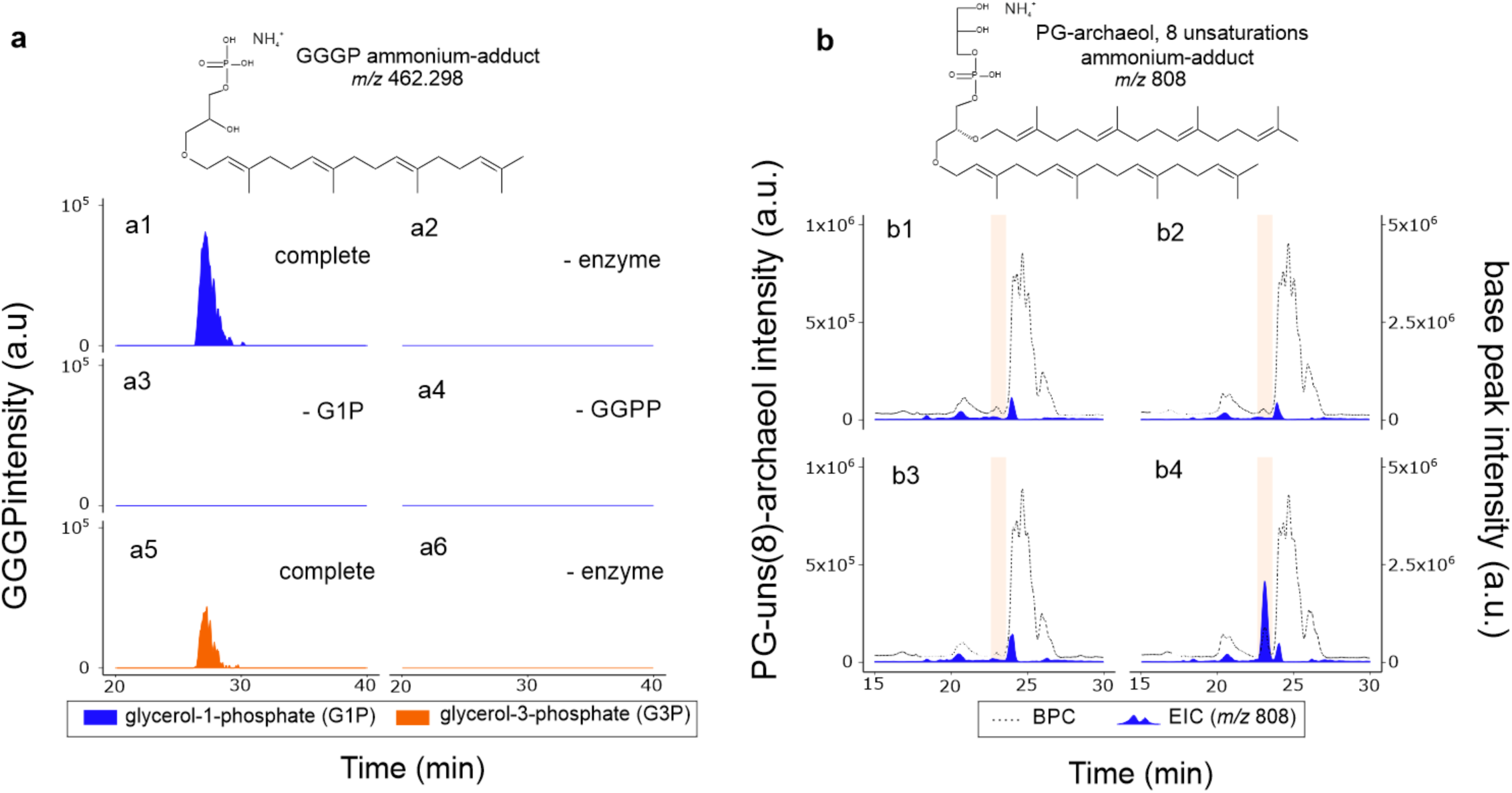
Biochemical verification of *Ca.* Cloacimonetes GGGP and DGGGP synthase activities. **a**, enzyme assay of purified GGGP synthase with G1P (**a1** to **a4**) or G3P (**a5** and **a6**). Extracted ion chromatogram, within 3 ppm mass accuracy, of [GGGP+NH_4_]^+^ (*m/z* 462.298) of complete enzymatic assay (**a1** and **a5**) or control assays lacking enzyme (**a2** and **a6**), glycerol-phosphate (**a3**) or geranylgeranyl-diphosphate, GGPP (**a4**). **b**, Production of the lipid PG-archaeol with 8 double bonds or unsaturations (PG-unsat(8)-archaeol) in *E. coli*. Extracted ion chromatogram (± 0.5, mass units, mu) of [PG-unsat(8)-archaeol+H]^+^ (*m/z* 808); blue filled area, left axis) or base peak chromatogram (dotted line, right axis) of optimized *E. coli* containing empty-vector (**b1**) or vector encoding GGGP synthase (**b2**), DGGGP synthase (**b3**) or both GGGP and DGGGP synthases (**b4**). Orange boxes spans the retention time of PG-unsat(8)-archaeol.

In addition to genes for GGGP and DGGGP synthase, other genes required for the synthesis of isoprenoidal archaeal lipids were also detected in the *Ca.* Cloacimonetes MAGs, including the genes for a complete bacterial isoprenoid MEP/DOXP pathway (Supplementary Table 3), as well as genes coding for acetyl-CoA C-acetyltransferase and hydroxymethylglutaryl-CoA synthase of the Mevalonate pathway (see Supplementary Table 3). Furthermore, two polyprenyl synthases were detected in the *Ca.* Cloacimonetes MAGs (see Supplementary Information, Supplementary Table 4). Finally, the *Ca.* Cloacimonetes MAGs also include digeranylgeranylglycerophospholipid reductases that are closely related to homologs found in other *Ca.* Cloacimonetes, as well as other members of the FCB group superphylum and related candidate phyla, which are related to homologs found in Euryarchaeota (see Supplementary Information). In summary, in addition to ether bond formation, these bacteria have the capacity to synthesize isoprenoid side chains and saturate them, characteristics that are fully in line with archaeal lipid membrane biosynthesis^1^.

Only two main genes of the archaeal lipid biosynthetic pathway were not detected in the *Ca.* Cloacimonetes MAGs. One of them is the CDP-archaeol synthase (CarS) forming the activated CDP-archaeol before the addition of the polar headgroups in combination with other specific archaeal enzymes^9^. However, it has been recently demonstrated that the bacterial CDP diacylglycerol synthase (CdsA) can replace the function of the archaeal CarS to generate CDP-archaeol. Subsequently, substrate promiscuity allows the bacterial phosphatidylglycerophosphate synthetase (pgsA) together with the phosphatidyl-glycerophosphatase (pgpA) to recognize CDP-archaeol and synthesize archaetidylglycerol^40^ as in the archaeal lipid biosynthetic pathway. All these bacterial enzymes are encoded by the four reported MAGs (Supplementary Table 2).

The second lacking gene is glycerol-1-phosphate dehydrogenase (egsA) coding for the enzyme enabling G1P biosynthesis (i.e. G1P-DH). Its bacterial homolog (araM), occurring in a few bacteria^41^, is also absent. This may seem enigmatic at first sight, as it would suggest that the presumed archaeal membrane lipids synthesized by *Ca.* Cloacimonetes would not have G1P as a glycerol phosphate backbone. Notably, there must be an alternative pathway for the formation of G1P, as G1P-archaeal membrane lipids were formed in a genetically engineered bacterial strain of *E. coli*, whose genome also did not contain egsA nor araM^11^. Considering this, it is possible that the archaeal-like membrane lipids synthesized by *Ca.* Cloacimonetes still have the archaeal G1P stereochemistry despite the lack of genes coding for G1PDH and AraM in their genomes. However, we cannot rule out that the presumed ‘archaeal’ lipids possess the bacterial G3P stereochemistry as promiscuity of GGGP synthase for G3P has been observed^11, 42, 43^.

Homology searches coupled to phylogenetic analysis indicated that the presence of archaeal lipid biosynthetic genes in bacterial genomes is not limited to *Ca.* Cloacimonetes from the Black Sea. Close GGGP and DGGGP synthase homologs are found together in other genomes of *Ca.* Cloacimonetes, as well as in other FCB group superphylum bacteria, genomes of related bacterial candidate phyla, and in one genome of *Candidatus* Parcubacteria of the Candidate Phyla Radiation^44^ (Fig. 3, Supplementary Tables 5 and 6). The phylogenetic topologies of both the GGGP and DGGGP synthases in bacteria are similar with respect to both branching of bacterial groups and sharing a close affiliation with GGGP and DGGGP synthases from the TACK group, in particular Crenarchaeota^32^, suggesting that the two genes share a similar evolutionary history (Fig. 3, Supplementary Information). Genomic co-localization of GGGP synthase and DGGGP synthase as observed in the four *Ca.* Cloacimonetes MAGs has previously only been seen in some Euryarchaeota, and within bacteria seems to be restricted to *Ca.* Cloacimonetes (Supplementary Information, Supplementary Table 7). Considering the basal placement of these genomes in the bacterial clade of both trees (Figure 3), we hypothesize that genomic co-localization of the genes may be the ancestral state.

**Figure 3.**
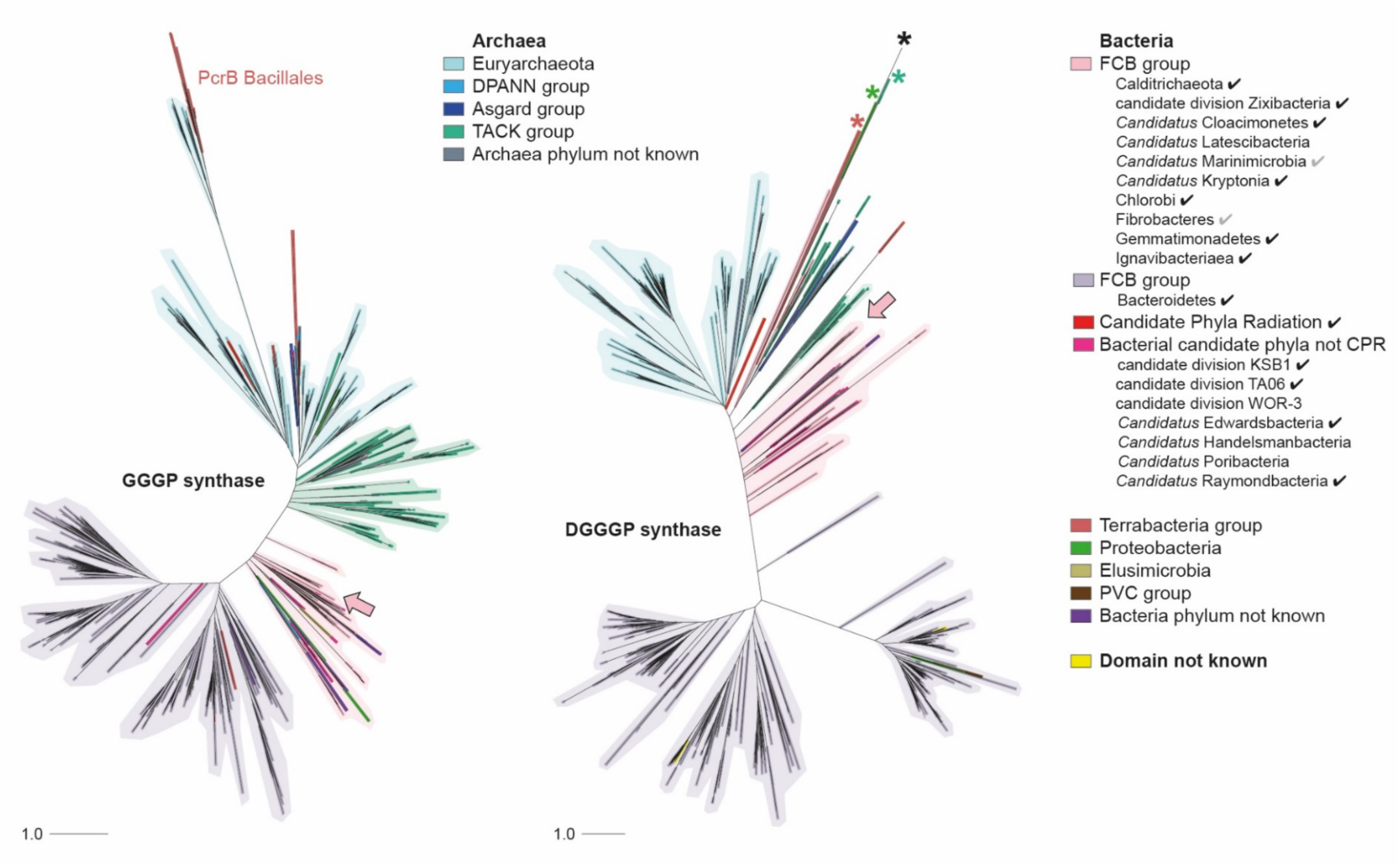
Phylogenetic tree of the GGGP and DGGGP synthase homologs detected across the tree of life. Search for homologs was performed with *Ca.* Cloacimonetes MAG sequences (arrows), in genomes from both cultures and environmental samples. Shadings illustrate the dominant group in the clade. Asterisks in the DGGGP synthase tree indicate known UbiA prenyltransferases that do not have DGGGP synthase activity. A checkmark in the legend marks bacterial groups with genomes that code for both GGGP and DGGGP synthase, a grey checkmark marks groups in which both genes are found but not in the same genome. The TACK group includes the Thaumarchaeota and the Crenarchaeota. Archaeal or Bacterial ‘phylum not known’: phylum is not known but the genome is annotated on a lower level, or sequence represents multiple groups. ‘Domain not known’: genomes for which no lineage was found on the PATRIC servers. Annotated trees with support values in Supplementary Figures 3 and 5.

The extended presence and close phylogenetic associations of the GGGP and DGGGP synthase homologs in the FCB group superphylum and related candidate phyla strongly supports the basal presence of these two enzymes in Bacteria, placing the origin well before the radiation of the superphylum. This implies that the capacity to synthesize ‘mixed membranes’ has clearly been stable in these bacteria over a long period of evolutionary time. Two evolutionary scenarios may explain our current observations. First, the contemporary presence of these enzymes in FCB group superphylum bacteria could be an evolutionary remnant of the ‘mixed membrane’ stage after LUCA and before the diversification of Bacteria. Second, if the ancestor of the FCB group superphylum contained a homochiral bacterial membrane like the other contemporary bacterial groups, the phylogenetic similarities of the genomically co-localized genes could also reflect ancient horizontal gene transfer (HGT) from an ancestral archaeal lineage into the FCB group ancestor (see Supplementary Information).

We have shown that members of the FCB group superphylum harbor a complete archaeal membrane lipid biosynthetic pathway. Our data strongly suggest that members of *Ca.* Cloacimonetes have the potential to synthesize a ‘mixed heterochiral membrane’, the first evidence of naturally occurring organisms with this ability. These organisms, as well as the rest of the FCB group superphylum, appear to be key in our understanding of the ‘lipid divide’, and are possible evolutionary remnants of the hypothetical ‘mixed membrane’ of LUCA. In any case, this discovery provides further support for the existence and thus feasibility of ‘mixed membranes’ in natural environments and over large evolutionary timescales, bridging the lipid divide.

## Methods

### Sampling

Suspended particulate matter (SPM) from 15 depths across the water column (50–2,000 m, Supplementary Table 8) was collected at sampling station 2 (N42°53.8’, E30°40.7, 2,107m depth) in the Black Sea western gyre during the Phoxy cruise 64PE371 (BS2013) on the 9^th^ and 10^th^ June 2013 on board the R/V Pelagia (Supplementary Table 8). In addition, SPM was also collected at 1,000 m, 1,500 m, and 1,980 m (BS2016) in the same location during the NESSC Black Sea cruise 64PE408 in 31^st^ January–2^nd^ February 2016 also on board of the R/V Pelagia. SPM was collected with McLane WTS-LV *in situ* pumps (McLane Laboratories Inc., Falmouth) on pre-combusted glass fiber (GF/F) filters with 142 mm diameter and 0.7 µm pore size (Pall Corporation, Washington) stored at −80°C. Water samples were also collected simultaneously at the same depths to perform physicochemical analysis (Supplementary Table 8).

### Nucleic acid extraction and 16S rRNA gene amplicon sequencing

DNA and RNA were extracted from sections of the 0.7 µm GF/F filters (1/8 filter from 50–130 m depth and ¼ from 170–2,000 m depth) with the RNA PowerSoil^®^ Total Isolation Kit plus the DNA elution accessory (Mo Bio Laboratories, Carlsbad, CA). RNA extracts were treated with DNAse and reverse-transcribed to cDNA using random nonamers as described previously^45^. The 16S rRNA gene amplicon sequencing and analysis was performed as described previously^46^. Taxonomy of the reads was assigned based on blast and the SILVA database version 123^47, 48^.

### Metagenome sequencing and assembly

Unamplified DNA extracts from the 15 SPM samples were used to prepare TruSeq nano libraries which were further sequenced with Illumina Miseq (5 samples multiplexed per lane) at Utrecht Sequencing Facility, generating 45 million 2×250 bp paired-end reads. Quality control was performed with FastQC v0.11.3 (https://www.bioinformatics.babraham.ac.uk/projects/fastqc/), and reads with uncalled bases and remaining TruSeq adapters were removed with Flexbar v2.5^49^, keeping the longer side of the read with the ‘--ae any’ flag. All reads were cross-assembled with SPAdes v3.8.0 in ‘--meta’ mode^50^, with read error correction turned on. BWA-MEM v0.7.12-r1039^51^ was used to map the forward and reverse reads from individual samples to the cross-assembled scaffolds.

### Scaffold binning and assessment of MAG quality

Scaffolds were binned into draft genome sequences based on coverage profile across samples and tetra-nucleotide frequency with MetaBAT v0.32.4^52^. The ‘--superspecific’ preset was used to minimize contamination. To increase sensitivity without losing specificity, MetaBAT was run with ensemble binning, which aims to combine highly similar bins (‘-B’ and ‘--pB’ were set to 20% and 50%, respectively). Quality of the MAGs was assessed based on absence and presence of lineage-specific marker gene sets after genome placement in a reference tree with CheckM v1.0.7^53^ in ‘--lineage_wf’ mode.

### Genome annotation and abundance estimation

MAGs were annotated with Prokka v1.11^54^ in ‘--metagenome’ mode. GenBank output files generated by Prokka were also annotated with the Rapid annotation using subsystem technology (RAST) pipeline v2.0^55^ (Supplementary Tables 9–12). MAG abundance was estimated from the shotgun metagenomics data by generating depth files per sample for all the scaffolds with SAMtools v.1.3.1^56^, using the mpileup utility with flags ‘-aa’ and ‘-A’ (count orphans) set. Average read coverage per nucleobase of a scaffold was calculated by dividing the sum of depth of all positions by the length of the scaffold with N’s removed. Coverage of a MAG was calculated likewise after concatenating the depth files of the scaffolds in that MAG. Average read coverage per nucleobase was normalized across samples by dividing by the total number of reads after quality control in the sample times 1,000,000. Normalized data is only shown in Figure 1b.

### Manual cleaning

The four *Ca.* Cloacimonetes MAGs were cleaned by plotting coverage across samples for all the scaffolds in the MAG, and manually removing those scaffolds that did not clearly have the same coverage profile as the majority (Supplementary Figure 1). NIOZ-UU3 seemed clean based on its coverage profile so none of its scaffolds were removed. Because the removed scaffolds do not contain marker genes, completeness and contamination estimates did not change (See Supplementary Table 1 for extended CheckM results of the four MAGs). Genome annotations and abundance estimations of MAGs were newly generated as described above after cleaning.

### Placement of MAGs in the FCB group superphylum tree

To further check the phylogenetic affiliation of the four MAGs in comparison with close relatives, Bacteroidetes/Chlorobi group genomes were downloaded from RefSeq^57^, and other FCB group genomes from GenBank^58^, both on February 8^th^, 2017 (Supplementary Table 13). For Ignavibacteriae and Chlorobi all the RefSeq representative genomes were downloaded, and for Bacteroidetes only the first ten representative genomes. We only included genomes that were estimated to be less than 10% contaminated as called by CheckM in ‘--lineage_wf mode’, and that contained at least 4 out of 43 phylogenetically informative marker genes in single copy that CheckM uses for bin placement^53^ (Supplementary Table 13). We realigned the 43 marker genes individually with Clustal Omega v1.2.3^59^. The genes were concatenated, gaps included if a gene was not present, and identical sequences removed (Supplementary Table 13). A maximum likelihood tree was inferred with RAxML v8.2.9^60^, with flags set to ‘-f a -m PROTCATLG-N 100-p 12345-x 12345’. In short, 100 rapid bootstrap searches were performed under the CAT approximation, and final trees were more thoroughly evaluated assuming a gamma distribution of rate heterogeneity for variable sites, with shape parameter α estimated. Amino acid substitution rates were based on the LG matrix. The best scoring tree was visualized with iTOL^61^.

We identified 16S rRNA genes in the FCB group genomes included in the tree with the CheckM ‘ssu_finder’ utility (Supplementary Table 13), resulting in 16S rRNA gene sequences for 20 *Ca.* Cloacimonetes genomes including NIOZ-UU1. We aligned these sequences with the *Ca.* Cloacimonetes amplicon sequences using MAFFT v7.394^62^, sliced out the amplicon region from the genome sequences, and removed identical sequences (Supplementary Table 13).

### Homology search of GGGP synthase and DGGGP synthase across the tree of life and gene tree construction

Predicted GGGP and DGGGP synthases from the four MAGs were queried with blastp^47^ against 110,421 annotated genomes available in the PATRIC genome database^63^ on November 29^th^, 2017. Blastp was run per genome (e-value <1e-10 and ≥70% query coverage) with a fixed database size of 20,000,000 to make e-values comparable across genomes. For GGGP synthase, we collected all hits and included the entire protein in the analysis. To exclude other prenyltransferases from the superfamily, for DGGGP synthase we only included hits that were annotated as ‘Digeranylgeranylglyceryl phosphate synthase (EC 2.5.1.42)’, and (‘similar to’) ‘(S)-2,3-di-O-geranylgeranylglyceryl phosphate synthase’, and hypothetical proteins with ≥90% query coverage. Again, entire proteins were included in the analysis. Moreover, we included a set of known GGGP and DGGGP synthases based on biochemical evidence and phylogenetic analyses^64^, as well as other non-DGGGP prenyltransferases as ‘outgroups’ (see Supplementary Information). The protein sequences were aligned after a first round of identical sequence removal with MAFFT v7.394^62^, using a maximum number of 1,000 iterative refinements and local pair alignment (L-INS-i). The sequence alignments were trimmed with trimAl v1.4.rev22^65^ in ‘-gappyout’ mode, followed by a second round of identical sequence removal. Final alignment lengths were 246 and 266 positions for GGGP synthase and DGGGP synthase, respectively. Model selection of nuclear models was performed with ModelFinder^66^ in IQ-TREE^67^ and maximum likelihood trees were constructed with the best-fit model chosen according to the Bayesian Information Criterion (LG+R10 for GGGP synthase and LG+F+R10 for DGGGP synthase), including 1,000 ultrafast bootstraps^68^ (random seed set to 12,345). For both trees, the consensus tree had a higher likelihood than the maximum likelihood tree found. Major clade separations were comparable between the maximum likelihood and consensus tree for both genes. Consensus trees were visualized in iTOL^61^.

### Co-localization of GGGP and DGGGP synthase in FCB group superphylum genomes

Predicted GGGP and DGGGP synthases from the four *Ca.* Cloacimonetes MAGs were queried with tblastn^47^ (e-value <1e^-5^ and ≥70% query coverage) against all downloaded GenBank/RefSeq FCB group superphylum genomes (above) to identify homologs. If hits were located on the same scaffold, the minimum base pair distance between GGGP and DGGGP synthase homologs was considered as a measure of co-localization.

### PCR amplification, sequencing and gene expression of specific genes detected in NIOZUU3

To experimentally assess the assembly accuracy of NIOZ-UU3, primers were designed to amplify and sequence the genes predicted to code for the GGGP, DGGGP, polyprenyl synthase and the bacterial marker gene predicted by CheckM (helicase PriA see Supplementary Information) from the BS2013 2,000 m sample. PCR products were gel purified (QIAquick gel purification kit, Qiagen), cloned in the TOPO-TA cloning^®^ kit from Invitrogen (Carlsbad, CA, USA). PCR targeting the coding genes of the GGGP synthase, DGGGP synthase, and polyprenyl synthase were tested with DNA and cDNA from the SPM samples recovered at 1,000 m and 2,000 m from the Black Sea 2013 campaign as a template.

### Construction of GGGPS and DGGGPS expression plasmids

Plasmids for *in vivo* lipid production were constructed by PCR amplifying the GGGP synthase ORF from pLVA01 and the plasmid pRSFDuet-1 (Novagen) using the primers GGGPS-RSFDuet-F/R and RSFDuet-GGGPS-F/R, respectively, and further recombined^69^ (Supplementary Information). A non-synonymous mutation (Met to Val, based on NIOZ-UU3) was corrected by site-directed mutagenesis using the primers GGGPS-SDM-F/R, resulting in pABW1. The DGGGP synthase ORF was amplified using the primers DGGGPS-F/R and cloned into pRSFDuet-1 using NcoI and BamHI to construct pABW2. pABW3 (encoding both GGGPS and DGGGPS) was constructed by re-amplifying DGGPS from pABW2 and cloning the amplicon as a NcoI-BamHI fragment into pABW1. A N-terminally 6His-tagged GGGP synthase overproduction plasmid (pABW4) was constructed by recombining^69^ the GGGP synthase ORF from pABW1 and pET-28a(+) (Novagene) amplified using the primers GGGPS-ET28-F/R and ET28-GGGPS-F/R, respectively. All inserts were verified to encode correct proteins by sequencing.

### Recombinant production, purification and *in vitro* enzyme assay of GGGP synthase

Purification and enzymatic assay of the NIOZ-UU3 GGGP synthase was based on the method of Jain et al.^70^. *E. coli* BL21(DE3) harboring pABW4 was cultured in 250 mL Lysogeny broth (LB) medium (37 °C, 200 rpm) and induced with 0.5 mM isopropyl β-D-1-thiogalactopyranoside (IPTG) at 0.6 OD_600nm_ for 4 h. Cells were centrifuged (3,500 rcf) and frozen. Subsequent steps were performed at room temperature, with samples and buffers kept on ice. Thawed cells were resuspended in ∼ 5 mL lysis buffer (50 mM Tris-HCl pH 7.5, 150 mM NaCl, 20 mM imidazole, 1 mg/mL lysozyme and a protease inhibitor (cOmplete™, EDTA-free; Roche, Basel) and sonicated to facilitate cell lysis. The lysate was cleared by centrifugation (20,000 rcf, 10 min), glycerol was added (10% final), and sample loaded onto a gravity column containing 1.5 mL of nickel-nitrilotriacetic acid (Ni-NTA) agarose beads (Qiagen, Venlo, NL) pre-equilibrated with protein buffer (50 mM Tris-HCl pH 7.5, 150 mM NaCl, 20 mM imidazole, 10% glycerol). Beads were washed with ∼ 30 column volumes of protein buffer, and initially eluted by a titration of imidazole with GGGP synthase protein eluting at 200 mM. Subsequent elutions were performed with 250 mM imidazole. The purity of the GGGP synthase protein was verified using 12% TGX^TM^ precast gels (Bio-Rad), stained with Bio-Safe™ Coomassie stain (Bio-Rad). The concentration of the purified protein was calculated based on the absorbance at 280 nm using a predicted extinction coefficient of 9,002 M^-1^ cm^-1^ (ExPASy ProtParam, averaged Cys reduced and cysteine form).

Enzymatic activity of the purified *Ca.* Cloacimonetes GGGP synthase was determined in an end-point assay using 0.1 µM GGGP synthase, 10 mM glycerol-1-phosphate (G1P) or glycerol-3-phosphate (G3P) and 100 µM geranylgeranylpyrophosphate in a reaction buffer consisting of 50 mM Tris-HCl (pH 7.5), 10 mM MgCl, and carryover amounts of imidazole (1 mM) and glycerol (0.5%). Reactions were incubated for 2 h at 37 °C in glass vials, extracted twice with 300 µL n-butanol (water saturated), pooled and stored at −20 °C. Analysis of the GGGP synthase enzyme assays were performed using Ultra High Pressure Liquid Chromatography – High Resolution Mass Spectrometry (UHPLC-HRMS) based on Sturt et al.^71^ with some modifications as detailed below. Pooled butanol extracts were evaporated under a stream of nitrogen, redissolved in 50 µL methanol:dichloromethane (1:1) and filtered (0.45 µm, regenerated cellulose). Analysis was performed using an Agilent 1290 Infinity I UHPLC, equipped with thermostatted auto-injector and column oven, coupled to a Q Exactive Orbitrap MS with Ion Max source with heated electrospray ionization (HESI) probe (Thermo Fisher Scientific, Waltham, MA). Injection volume was 1 µl (out of 50 µl). Separation was achieved on a YMC-Triart Diol-HILIC column (250 × 2.0 mm, 1.9 µm particles, pore size 12 nm; YMC Co., Ltd, Kyoto, Japan) maintained at 30 °C. The following elution program was used with a flow rate of 0.2 mL min^-1^: 100% A for 5 min, followed by a linear gradient to 66% A: 34% B in 20 min, maintained for 15 min, followed by a linear gradient to 40% A: 60% B in 15 min, followed by a linear gradient to 30%A:70%B in 10 min, where A = hexane/2-propanol/formic acid/14.8 M NH_3aq_ (79:20:0.12:0.04 [volume in volume in volume in volume, v/v/v/v]) and B = 2-propanol/water/formic acid/14.8 M NH_3aq_ (88:10:0.12:0.04 [v/v/v/v]). HESI settings were as follows: sheath gas (N_2_) pressure 35 (arbitrary units), auxiliary gas (N_2_) pressure 10 (arbitrary units), auxiliary gas (N2) T 50 °C, sweep gas (N_2_) pressure 10 (arbitrary units), spray voltage 4.0 kV (positive ion ESI), capillary temperature 275 °C, S-Lens 70 V. Lipids were analyzed with a mass range of *m/z* 350–2,000 (resolving power 70,000) followed by data dependent MS^2^ (resolving power 17,500), in which the ten most abundant masses in the mass spectrum (with the exclusion of isotope peaks) were fragmented successively (stepped normalized collision energy 15, 22.5, 30; isolation window 1.0 *m/z*). An inclusion list was used with a mass tolerance of 3 ppm, targeting the ammoniated molecule [C_46_H_79_O_8_P+NH_4_]^+^ of GGGP at *m/z* 462.2979. The Q Exactive was calibrated within a mass accuracy range of 1 ppm using the Thermo Scientific Pierce LTQ Velos ESI Positive Ion Calibration Solution (containing a mixture of caffeine, MRFA, Ultramark 1621, and *N*-butylamine in an acetonitrilemethanol-acetic solution). Identification of GGGP was aided by the analysis of 1-O-octadecyl-2-hydroxy-*sn*-glycero-3-phosphate (C_18_-LPA; Avanti Polar Lipids, Inc. Alabama, USA), a structural analogue of GGGP, which has a C_18_ carbon chain attached to the glycerol backbone instead of the geranylgeranyl carbon chain present in GGGP. C_18_-LPA showed similar chromatographic and mass spectral behavior to GGGP.

### Co-expression of GGGP and DGGGP synthases in *E. coli*

*E. coli* C43(DE3)^72^ harboring GGPP synthase (*crtE*) and G1PDH (*araM*) on plasmid pMS148^11^ was used for expression of the GGGP and DGGGP synthases (encoded on plasmids pABW1, −2 and −3). Cells growing exponentially in LB medium were diluted into magnesium-supplemented terrific broth medium^73^ (Mg-TB; 1.2% Tryptone, 2.4% Yeast Extract, 0.4% glycerol, 2 mM MgSO_4_, 0.23% KH_2_PO_4_ and 1.25% K_2_HPO_4_) to 0.01 OD_600nm_ induced with 0.4 mM IPTG and incubated at 37 °C (200 rpm) for 16 h. Cells (10 mL culture normalized to 1.9 OD_600nm_) were harvested by centrifugation (4,000 rcf, 10min) and washed twice with 0.85% NaCl, lyophilized and then stored at −80°C. Analysis for production of archaeal-like lipids in *E. coli* was performed by extracting intact polar lipids (IPLs) by a modified Bligh-Dyer extraction^74^ that was analyzed according to^75^ with some modifications: lyophilized cells were extracted thrice with BDE solvent mixture (2:1:0.8 methanol:dichloromethane (DCM):potassium phosphate buffer (50 mM, pH 7) aided by sonication and centrifugation. The extracts were pooled and solvent ratios adjusted to 1:1:0.9, vigorously mixed, centrifuged and the lower DCM phase transferred. The upper fraction was re-extracted twice with DCM, and the pooled extract was evaporated under a stream of nitrogen and stored dry at −20 °C until analysis. For analysis, samples were dissolved in 200 µl hexane:isopropanol:H_2_O (718:271:10) and filtered (0.45 µm, regenerated cellulose. Analysis was performed on an Agilent 1200 series LC (Agilent, San Jose, CA), equipped with thermostatted auto-injector and column oven, coupled to a Thermo LTQ XL linear ion trap with Ion Max source with electrospray ionization (ESI) probe (Thermo Scientific, Waltham, MA), was used. Separation was achieved on a YMC-Pack-Diol-120-NP column (250 × 2.1 µm, 5 µm particles; YMC Co., Ltd, Japan) maintained at 30 °C. Elution program and ESI settings are described in^75^. The lipid extract was analyzed by an MS routine where a positive ion scan (*m/z* 400–2,000) was followed by a data dependent MS^2^ experiment where the base peak of the mass spectrum was fragmented (normalized collision energy (NCE) 25, isolation width 5.0, activation Q 0.175). This was followed by a data dependent MS^3^ experiment where the base peak of the MS^2^ spectrum was fragmented under identical fragmentation conditions. This process was repeated on the 2^nd^ to 4^th^ most abundant ions of the initial mass spectrum.

## Acknowledgements.

We thank Melvin Siliakus for providing several of the plasmid constructs and for useful suggestions to improve the manuscript. We are also thankful to Julian Vosseberg, John van Dam, Anja Spang, and Jan de Leeuw for suggestions and constructive discussions. We acknowledge the Utrecht Sequencing Facility (USF), which is partially subsidized by Hubrecht Institute, Utrecht University, and UMC Utrecht, for the sequencing data and service. We thank Elda Panoto, Maartje Brouwer and Michel Koenen for providing technical support. We acknowledge the crew and scientists of the R/V Pelagia cruises 64PE371 (chief scientist Gert-Jan Reichart) and 64PE408 (chief scientist Marcel van der Meer). J.S.S.D. received funding from the European Research Council (ERC) under the European Union’s Horizon 2020 research and innovation program (grant agreement n° 694569—MICROLIPIDS). L.V. and J.S.S.D. receive funding from the Soehngen Institute for Anaerobic Microbiology (SIAM) through a Gravitation Grant (024.002.002) from the Dutch Ministry of Education, Culture and Science (OCW). B.E.D. is supported by the Netherlands Organization for Scientific Research (NWO) Vidi grant 864.14.004.

## Author contributions

L.V., B.E.D., and J.S.S.D. conceived the study. L.V., F.A.B.v.M., and B.E.D. analyzed environmental sequencing data and performed phylogenetic analyses. A.B.W. performed studies of the recombinant proteins. A.B.W. and E.C.H. analyzed enzymatic assays and cells using LC-MS. L.V., F.A.B.v.M, and J.S.S.D. wrote, and all authors edited and approved the manuscript.

## Data availability and author information

The 16S rRNA gene amplicon reads (raw data) have been deposited in the NCBI Sequence Read Archive (SRA) under BioProject number PRJNA423140. The *Ca.* Cloacimonetes MAGs are deposited in IMG under the following IMG accession IDs: 134200 (NIOZ-UU1), 134201 (NIOZ-UU2), 134202 (NIOZ-UU3), 151202 (NIOZ-UU4). The authors declare no competing financial interests. Correspondence and requests should be addressed to L.V. and F.A.B.v.M.

## Supplementary information

Supplementary results and discussion, Supplementary Figures and Supplementary Tables are available for this paper.

